# Estrogen regulates myogenic tone in hippocampal arterioles by enhanced basal release of nitric oxide and endothelial SK_Ca_ channel activity

**DOI:** 10.1101/2023.08.15.553442

**Authors:** Danielle A. Jeffrey, Abigail Russell, Mayra Bueno Guerrero, Jackson T. Fontaine, Phinea Romero, Amanda C. Rosehart, Fabrice Dabertrand

**Author notes:** Correspondence to: Fabrice Dabertrand, Ph.D., Email address, Postal address:, Research Complex 2 (Rm 7120 - Mailstop B112), 12700 E 19th Avenue, Aurora CO 80045, USA.

## Abstract

Arteries and arterioles exhibit myogenic tone, a partially constricted state that allows further constriction or dilation in response to moment-to-moment fluctuations in blood pressure. The vascular endothelium that lines the internal surface of all blood vessels controls a wide variety of essential functions, including the contractility of the adjacent smooth muscle cells by providing a tonic vasodilatory influence. Studies conducted on large (pial) arteries on the surface of the brain have shown that estrogen lowers myogenic tone in female mice by enhancing nitric oxide (NO) release from the endothelium, however, whether this difference extends to the intracerebral microcirculation remains ambiguous. The existing incomplete picture of sex differences in cerebrovascular physiology combined with a deficiency in treatments that fully restore cognitive function after cerebrovascular accidents places heavy emphasis on the necessity to investigate myogenic tone regulation in the microcirculation from both male and female mice. We hypothesized that sex-linked hormone regulation of myogenic tone extends its influence on the microcirculation level, and sought to characterize it in isolated arterioles from the hippocampus, a major cognitive brain area. Using diameter measurements both *in vivo* (acute cranial window vascular diameter) and *ex vivo* (pressure myography experiments), we measured lower myogenic tone responses in hippocampal arterioles from female than male mice. By using a combined surgical and pharmacological approach, we found myogenic tone in ovariectomized (OVX) female mice matches that of males, as well as in endothelium-denuded arterioles. Interestingly, eNOS inhibition induced a larger constriction in female arterioles but only partially abolished the difference in tone. We identified that the remnant difference was mediated by a higher activity and expression of the small-conductance Ca^2+^-sensitive K^+^ (SK) channels. Collectively, these data indicate that eNOS and SK channels exert greater vasodilatory influence over myogenic tone in female mice at physiological pressures.

## Introduction

The mammalian cardiovascular system is sexually dimorphic. One consequence is that cardiovascular diseases develop more swiftly in men relative to women before menopause, while remaining the leading cause of death globally^1^. The major female sex hormone estrogen as long been recognized as a key element in this difference^2^. Estrogen, especially its most common circulating form 17β-estradiol (E2), is cardioprotective through a plethora of effects including, but not limited to, anti-inflammatory and antioxidant properties, plasma cholesterol regulation, and blood pressure lowering through vasodilation^3^. This last effect has received considerable research attention as hypertension is still the major, modifiable, risk factor for cardiovascular diseases^4^. Focusing on large blood vessels like coronary^5^, carotid^6^, or brain surface (pial) arteries^7,8^ early studies have provided unequivocal evidence that estrogen enhances the production of nitric oxide (NO), a potent vasodilator produced by the endothelial cells lining the lumen of all blood vessels^9,10^. However, it is important to note that the microcirculation has largely been unexplored in the context of mechanistic sex differences, especially within the brain—an exquisite target of cardiovascular diseases.

The vascular endothelium is uniquely positioned to respond to circulating hormones. It controls a wide variety of essential functions, beginning with the contractility of the adjacent smooth muscle cells (SMCs). Resistance arteries and arterioles exist in a partially constricted state called myogenic tone^11^. A raise in blood pressure activates contractile mechanisms increasing myogenic tone, which reduces vascular diameter, thus increasing vascular resistance. Conversely, a decline in blood pressure decreases vascular resistance by relaxing SMCs^12^. Myogenic tone is thus crucial to blood flow regulation over a wide range of arterial pressure. In the brain, it not only protects the capillary bed against disruptive blood pressure by supporting autoregulation, but it also creates the vasodilatory reserve necessary for functional hyperemia^13^. Mechanistically, the myogenic response is initiated by a gradated pressure-dependent depolarization of the SMC plasma membrane, which increases the open probability of the voltage-dependent Ca^2+^ channels (VDCC), increasing Ca^2+^ entry and promoting vasoconstriction^11,14,15^. In both brain pial arteries and parenchymal arterioles, basal NO release by the endothelium acts as a constitutive brake on pressure-induced depolarization and constriction by relaxing SMCs^16^. This is well demonstrated by the vasoconstriction induced by inhibitors of NO synthase (NOS) such as Nω-Nitro-L-arginine methyl ester hydrochloride (L-NAME) in both vascular beds^17^. In the endothelium, estrogen has been found to enhance both expression and activity of eNOS, the endothelial form of the enzyme^18,19^. Yet, profound differences exist between the pial and parenchymal endothelium vasodilatory influence, especially in the tonic hyperpolarizing input from endothelial K^+^ channels.

Activation of endothelial intermediate (IK) or small (SK) conductance Ca^2+^-activated K^+^ channels lead to both endothelium and SMCs hyperpolarization, which is then referred to as “endothelium-dependent hyperpolarization” (EDH) and is communicated through gap junctions via the myoendothelial projections to the adjacent SMCs^20^. Inversely, SMC membrane potential hyperpolarization deactivates VDCC, leading to vasodilation. In the pial and systemic arteries, EDH in response to endothelial agonist is blocked by IK and SK channels inhibitors. However, these blockers have little effect by themselves on the baseline diameter, revealing minimal IK/SK channel activity under basal conditions^21,22^. On the other hand, studies on intracerebral arterioles from our group and others have established that endothelial IK and SK channels have a significant tonic activity, continuously opposing pressure-induced constriction, even in the absence of exogenous agonist^17,23,24^. Remarkably, the combined IK/SK current density in parenchymal arteriolar ECs^24^ is more than 8-fold higher than those measured in ECs from large arteries^25-27^. This high current density coupled to a single layer of SMCs further shapes the high tonic control of parenchymal arteriolar tone by SK and IK channels. Importantly, estrogen has been proposed to regulate the expression of *Kcnn3*, the gene encoding SK channels both in *in vitro* cell models and mesenteric arteries^28,29^ however the brain microcirculation has been largely neglected. Thus, further characterization of the effects of estrogen and SK channel function and expression in the brain is warranted.

Brain blood flow is precisely and dynamically regulated by parenchymal arteriole diameters^30-32^ and cardiovascular diseases are recognized as major risk factors for vascular cognitive impairment, the major cause of dementia after Alzheimer’s disease^33^. Precise understanding of the regulation of parenchymal arteriole diameter, and the sex differences underlying it, is then crucial for healthy brain aging. Here we tested the hypothesis that estrogen decreases myogenic tone in mouse cerebral female arterioles by a unique combination of eNOS and SK channel increased activity. We focused on hippocampal parenchymal arterioles (HiPAs) as a cognitive-centric region known to be vulnerable to vascular dementia.

## Materials and Methods

### Animals

All experimental protocols used in this study were approved by the Institutional Animal Care and Use Committee (IACUC) of the University of Colorado, Anschutz Medical Campus. Adult (2–4 months old) male and female C57/BL6J mice (Jackson Laboratories, United States), were housed on a 12-h light:dark cycle with environmental enrichment and free access to food and water. All mice were euthanized by intraperitoneal injection of sodium pentobarbital (100 mg/kg) followed by rapid decapitation.

### Myography

Hippocampal parenchymal arterioles (HiPAs) were obtained as previously described^34,35^. Briefly, after euthanasia, the brain was removed and placed into chilled (4°C). MOPS-buffered saline (composition:135 mM NaCl, 5 mM KCl, 1 mM KH_2_PO_4_, 1 mM MgSO_4_, 2.5 mM CaCl_2_, 5 mM glucose, 3 mM MOPS, 0.02 mM EDTA, 2 mM pyruvate, 10 mg/mL bovine serum albumin, pH 7.3 at 4 °C). Arterioles that emanate from the anterior, middle, and posterior hippocampal arteries were dissected free of the surrounding tissue, cannulated on borosilicate glass micropipettes (with one end occluded) in an organ chamber (University of Vermont Instrumentation and Model Facility), and pressurized using an arteriography system (Living Systems Instrumentation, Inc., St. Albans, VT, USA). Vessel internal diameter was continuously monitored using a CCD camera and edge-detection software (IonOptix, Westwood, MA, USA). HiPAs were perfused (4 mL/min) with prewarmed (36.5°C ± 1°C), gassed (5% CO2, 20 % O2, 75 % N2) artificial cerebrospinal fluid (aCSF) for at least 30 min. The composition of aCSF was 125 mM NaCl, 3 mM KCl, 26 mM NaHCO3, 1.25 mM NaH_2_PO_4_, 1 mM MgCl_2_, 4 mM glucose, 2 mM CaCl_2_, pH 7.3 at room temperature with gas aeration. Passive diameter was obtained in nominally Ca^2+-^free aCSF (0 mM [Ca^2+^]_o_ with 5 mM EGTA). Arteriolar tone was calculated with the following equation: [(passive diameter – active diameter)/(passive diameter)] × 100. Only viable HiPAs, defined as those that developed pressure-induced myogenic tone greater than 15 % at 40 mmHg, were used in the experiments. Changes in arterial diameter were calculated as the percentage change from base-line using the following equation [(change in diameter)/(initial diameter)] × 100.

### Acute cranial windows *in vivo*

Using a 150 ul retro-orbital injection, 3 mg/ml of tetramethylrhodamine (TRITC)-dextran was administered to mice under isofluorane (5% induction, 2% maintenance) anesthesia. An acute cranial window was then implanted as previously described^32,36^. Briefly, after TRITC-dextran injection, the skull was revealed and a cranial window above the somatosensory cortex that is roughly 2 mm in diameter was drilled. A custom stainless steel head place with a corresponding 2 mm hole was then placed over the exposed brain and secured to the skull using a mixture of superglue and dental cement. Throughout the surgery isoflurane anesthesia was weaned off while α-chloralose (50 mg/kg) and urethane (750 mg/kg) was administered via intraperitoneal injection. Mice were monitored throughout the surgery with a rectal thermometer and kept at 37°C with an electric heating pad.

### Endothelial denudation

Hippocampal microvasculature was isolated and cannulated on one end. Then a syringe with an air bubble was used to mechanically denude the endothelial cells as previously described^37^. Proper denudation was verified with application of the SK and SK channels agonist NS309 (1 μM), and only microvasculature that did not response to the endothelial vasodilator were used. Failed denuded HiPAs (absence of myogenic tone or dilatory response to NS309 application) were not included in this study.

### Immunohistochemistry (IHC)

Hippocampal microvasculature was isolated as previously described (Rosehart *JoVE* 2019; Fontaine *MAD* 2020). 48 hours prior to isolation, SLYGARD coated coverslips were plated and allowed to cure. Microvasculature was then cannulated and tied on one end and filled with 0.1% agrose in aCSF. This allowed the microvasculature to expand and be manipulated to appreciate all the arteriolar, transitional, and capillary segments on the SLYGARD coated coverslip which they adhered to (workflow displayed in Figure 5A). Microvasculature was then fixed in 10% buffered formalin phosphate for 45 minutes. Fixed vasculature was then washed 3X with phosphate buffered saline (PBS; 137 mM NaCl, 2.7 mM KCl, 8 mM Na_2_HPO_4_, and 2 mM KH_2_PO). The vasculature was then blocked and permeabilized in blocking buffer (2% BSA, 0.1% NP-40, in PBS). Antibodies were purchased from Abcam unless stated otherwise. The vasculature was incubated overnight in primary antibodies (all primary antibodies were incubated in 1:200 dilutions: SK Cat no. AB2200865, IK Fisher Scientific Cat no. MA5-27547, and CD31 Cat no. AB305267) at 4°C on a rocker. Slides were washed for 3X in PBS followed by secondary antibody incubation (all secondary antibodies were incubated in 1:200 dilutions: anti-mouse Fisher Scientific Cat no. PIA32727, anti-rabbit Cy5 Cat no. AB6564) at RT on a rocker for one hour. Slides were mounted with ProLong Diamond Antifade with DAPI (Cat no. P36966) and sealed with a coverslip and allowed to set overnight (at 4°C) before imaging. IHC secondary controls were utilized for all non-conjugated antibodies where only the secondary antibody was applied to indicate nonspecific binding.

### Ovariectomy (OVX)

Surgery was performed in 2–4 months old female mice. Surgical anesthesia was induced via chamber induction with 3% isoflurane and maintained with 1-2% isoflurane via nose cone in O_2_-enriched air. Surgical sites and the skin were prepared with three rounds of alternating betadine and 70% alcohol after the hair was shaved. Once anesthesia was induced, the first dose of analgesia was provided, meloxicam 2 mg/kg subcutaneous (SC). Under isoflurane anesthesia, a single skin incision (10 mm) was made and moved over to each side of the flank, where the muscles overlying the ovary were separated. The ovarian artery and vein were then ligated (5-0 or 6-0 absorbable suture) and the ovary was removed. Absorbable sutures were used to close the abdominal muscles. Bupivacaine (1-2mg/kg, SC in saline vehicle) was applied at each incision site (each flank and the midsagittal skin). The skin was then closed with skin staples. A single incision was sufficient to remove both ovaries, and staples were removed 7-14 days after surgery. Mice were given heat support in static caging before being returned to the housing room. Postoperative analgesia was provided by administration of meloxicam (2 mg/kg SC in saline vehicle) administered at the time of surgery and daily for 1-3 days after surgery. Animals were monitored for body weight, wound appearance, overall physical appearance, and activity levels to ensure the health of the animal during the first week of recovery. Microvasculature was then harvested as described above.

### Silastic implant

Subcutaneous hormone implantation was performed in 2–4 months old female mice following OVX. The silastic implant contained either saline control or estrogen provided a dose of 0.05-0.1mg/kg over the lifespan of the implant (about 3 weeks). OVX and hormone subcutaneous implants were performed under one anesthesia to limit the isoflurane exposure, then *ex vivo* studies were performed 2-3 weeks later. This period allows endogenous hormone levels to fall and exogenous levels to come to steady-state level (<u>Mannino</u> *J Pain* 2005; Ström *Scan J Clin Lab Invest* 2008; Ström *JoVE* 2012). A small incision (5 mm) was made beneath the base of the neck and blunt hemostats were used to make a small cavity for the implant. A silastic implant (made in-house, 1cm long silastic tubing filled with 2mg estradiol or saline vehicle) was placed in the cavity and bupivacaine was administered at the incision site (1-2mg/kg, SC in saline). The incision was then closed with skin staples. Mice were given heat support in static caging before being returned to the housing room. Postoperative analgesia was provided by administration of meloxicam (2 mg/kg SC in saline vehicle) administered at the start of surgery and daily for 1-3 days after surgery. Animals were monitored for body weight, wound appearance, overall physical appearance, and activity levels to ensure the health of the animal during the first week of recovery. Microvasculature was then harvested as described above.

### Reagents

Drug compounds were mixed to their respective molarities and infused with aliquots of aCSF before they were added to the chamber bath. Effects of drug exposure on vessel diameter were recorded using measurements described above. Endothelial function was tested by assessing the vasodilator response to IK and SK channels agonist NS309 at 1 μM. HiPAs not responding to NS309, typically with a near max vasodilation, were discarded from the study. NS309 (Cat no. 3895) and Paxilline (Cat no. 2006) was purchased from Tocris Bioscience (USA), L-NAME (Cat no. N5751), Apamin (Cat no. A95459), Charybdotoxin (Cat no. C7802) and all other chemicals and reagents were obtained from Sigma-Aldrich (USA).

### Fluorescent image processing, analysis, and statistics

*In vivo* cranial window two photon laser scanning microscopy (TPLSM) was performed using a Bruker Ultima Investigator multiphoton microscope using Prairie View software as previously described^36^. Parenchymal arterioles were identified by the direction of the red blood cells flowing into the brain. IHC confocal microscopy was captured using a Nikon Eclipse Ti2 with 0.22 um step sizes to create a Z-stack and appreciate the three-dimensional structure of the microvasculature. All images were blinded prior to uploading in Imaris software for analysis. Quantitative analysis of immunohistochemistry Z-stack images was performed using Imaris (Version 10.0) microscopy image analysis software. Gaussian filter was applied to all images to consistently reduce noise in Imaris. LabKit (FIJI plugin) machine learning was trained to detect fluorescence and identify puncta and create surfaces for each respective antibody for IHC or for network creation using TRIT-C as a marker for *in vivo* network analysis. The same trained algorithm was used for all Z-stacks for consistency in both IHC and cranial window *in vivo* studies. Each mouse imaged for IHC had 9 regions of interest (ROIs) of 15 μm in diameter) that were picked blindly. The volumes of each puncta within an ROI were summed giving a total volume of fluorescence per ROI for each antibody. CD31 was used as an endothelial cell marker (Figure 5B). Supplementary Figure 2 shows our Imaris quantitative workflow for IHC surface creation. For *in vivo* studies post network creation in Imaris, the average vessel diameter was calculated for 3 different regions in 4 independent mice from the same X/Y/Z dimensions both basal and post zero calcium aCSF application. Figure 1A shows the network creation filament using Imaris. All statistical tests were performed in GraphPad Prism 9. The statistical workflow is that all data were checked for normality (using Shapiro-Wilk, while frequency histogram and skewness were observed) and, if normally distributed, an unpaired t-test was used for groups of 2, while an ANOVA multiple comparisons was applied post hoc for groups larger than 2. Data in figures and text are presented as means ± standard error of the mean (SEM). All statistical methods are indicated in figure legends with corresponding independent numbers.

**Figure 1:**
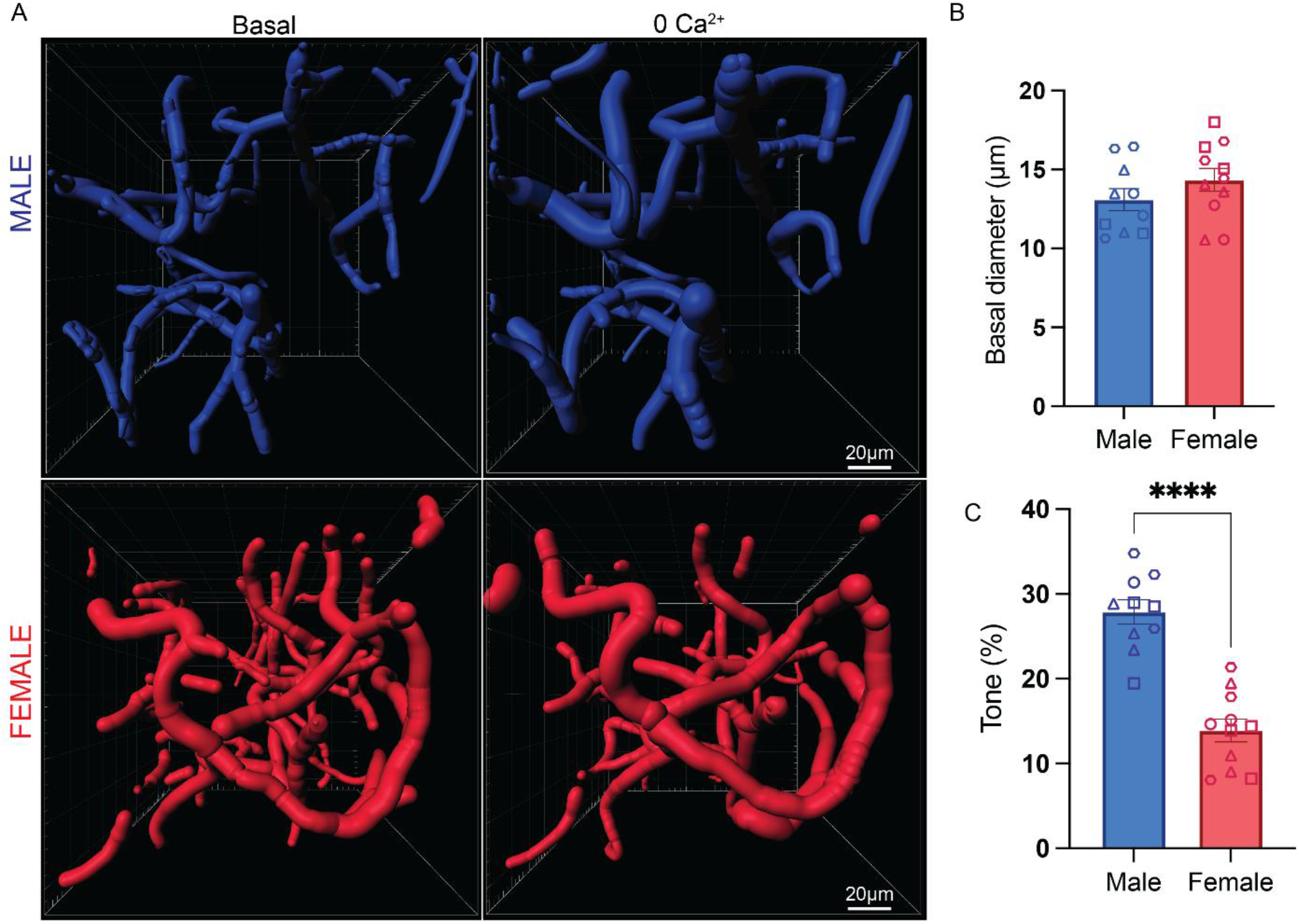
Cortical arterioles from female mice exhibit lower myogenic tone compared to male *in vivo*. (A) Superior view of vascular network Z-stack: Imaris software unbiased 3D reconstruction of cortical microcirculation visualized by two-photon microscopy following intravenous injection of TRITC-dextran. Top panels indicate males, bottom females, left panels show vessel diameters in basal conditions and right panels in absence of Ca^2+^. (B) Summary data showing average arteriole diameter. (C) Summary data showing significant differences in myogenic tone between males and females. Unpaired t test revealed statistically significant differences between male and female *in vivo* myogenic tone. **** indicates p value < 0.0001, n = 2-3 arteriole network subsections from 4 mice, symbols correlate with each individual animal (C/D).

## Results

### In Vivo Network Analysis Reveals Diminished Myogenic Tone in Female Mouse Parenchymal Arterioles Compared to Male

To compare the level of myogenic tone *in vivo* between male and female intracerebral arterioles, we first acquired cortical Z-stacks through a cranial window using two-photon laser-scanning microscopy in anesthetized mice. The parenchymal microcirculation was visualized by intravenous injection of tetramethylrhodamine (TRITC)-labeled dextran, and the skeletonization process to recreate the vascular network was achieved with Imaris software (Fig. 1A). We focused on vessels with an internal diameter of 10 μm and above to exclude capillaries from the analysis. Unbiased computational analysis with blinded images revealed no statistical difference in the diameter of male and female arterioles with an average of 13.10 ± 0.70 μm for the males and 14.14 ± 0.73 μm for the females (Fi. 1B). We then induced arteriolar maximal dilation by applying artificial cerebrospinal fluid (aCSF) depleted of Ca^2+^ over the cranial window (Fig 1A) and calculated the amount of myogenic tone. We found that myogenic tone in male arterioles was nearly twice as much as in female arterioles, 27.89 ± 1.42% vs. 13.90 ± 1.36%, respectively (Fig. 1C). This difference was highly significant and suggests that sex differences exist in the regulation of the myogenic tone of intracerebral arterioles.

### Hippocampal Parenchymal Arterioles (HiPAs) From Female Mice Exhibit Diminished Constriction to Pressure in the Physiological Range

To precisely compare myogenic tone between male and female, we measured changes in the diameter of isolated and pressurized parenchymal arterioles in response to increases in luminal pressure. We focused on the hippocampus (Fig. 2A) due to its susceptibility to pathological demise as it is the main brain region involved in learning and memory^38^. 20 mmHg of luminal pressure constricted HiPAs from male and female mice to the same extend ∼15% (Fig. 2B and C). However, at 40 mmHg and higher, arterioles from female mice constricted significantly less than arterioles from male mice. At 40 mmHg, the estimated pressure experienced *in vivo* by cerebral arterioles of this size^39^, female HiPAs constricted 12% less than male HiPAs (Fig. 2 D). Females had an average of 22.80 ± 0.96% tone at 40 mmHg while males had 34.74 ± 2.13% tone at 40 mmHg. This was significantly different per Bonferroni’s multiple comparisons test, p = 0.0008. The difference between male and female myogenic tone was significant at 40 mmHg, 60 mmHg, and 80 mmHg with response to pressure being consistently lower in females HiPAs (Figure 2D). These results confirm *ex vivo* sexual dimorphism in myogenic responses at various physiological and above pressures in hippocampal arterioles.

**Figure 2:**
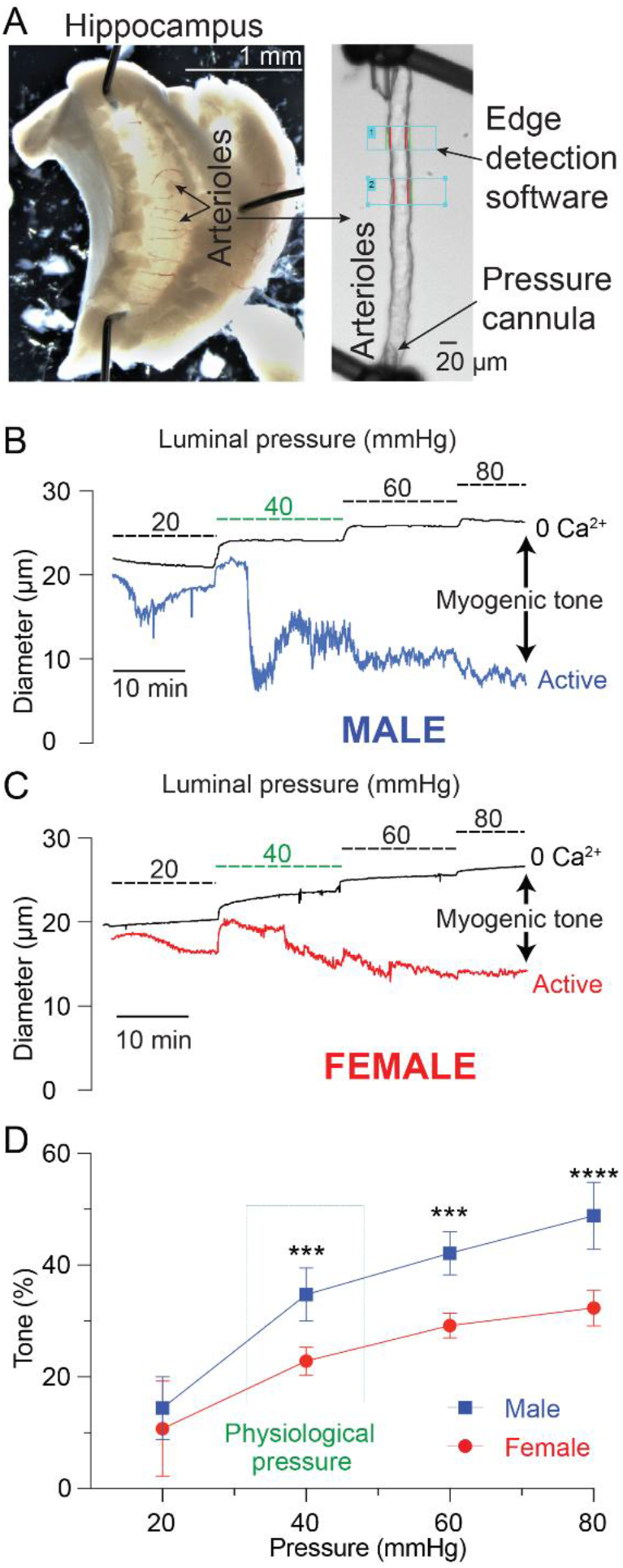
Arterioles from female hippocampi exhibit reduced levels of myogenic tone compared to male mice. (A) Micrograph of hippocampal arterioles in parenchyma (left panel) and isolated (right panel). (B) Representative traces of real-time diameter fluctuations of range of physiological pressures in myography experiments from male and (C) female mice. Pressure myography experiment traces track lumen diameter fluctuations over time (active) and are plotted next to traces in 0 Ca^2+^ aCSF (passive). (D) Measured myogenic tone over range of physiological blood pressures. Myogenic tone is calculated using the following equation [(passive diameter – active diameter)/(passive diameter)] × 100. Physiological pressure (40 mmHg) is noted. Post hoc bonferroni’s multiple comparison test indicated statistical significance. *** indicates p < 0.001, **** indicates p < 0.0001; n = 5 and 7 mice for males and females, respectively.

### The Diminished Myogenic Tone in HiPAs from female Mice Is Endothelium Dependent, But Only Partially Attributable to Increased NO Production

Previous studies conducted on large brain surface (pial) arteries revealed that lower myogenic tone in female mice is mediated by increased basal activity of eNOS^7,8^. We then compared the myogenic tone responses of HiPAs in which the endothelial function has been disrupted by passing an air bubble through the lumen^37^. Pressure-induced constriction was virtually identical between male and female endothelium-denuded HiPAs, validating a major role for the endothelium in generating this sexual dimorphism (Figure 3A-B). To specifically investigate the contribution of NO in lower female myogenic responses, we exposed arterioles pressurized at 40 mmHg to 100 μM of the NOS inhibitor L-NAME. HiPAs from both females and males constricted in response to L-NAME, indicating a constitutive eNOS activity. Consistent from previous findings ^7^, L-NAME-induced constriction was 17% higher in the female HiPAs (46.11 ± 4.72%), reflecting higher eNOS activity compared to males (29.33 ± 3.42%) (Fig 3C-D).

**Figure 3:**
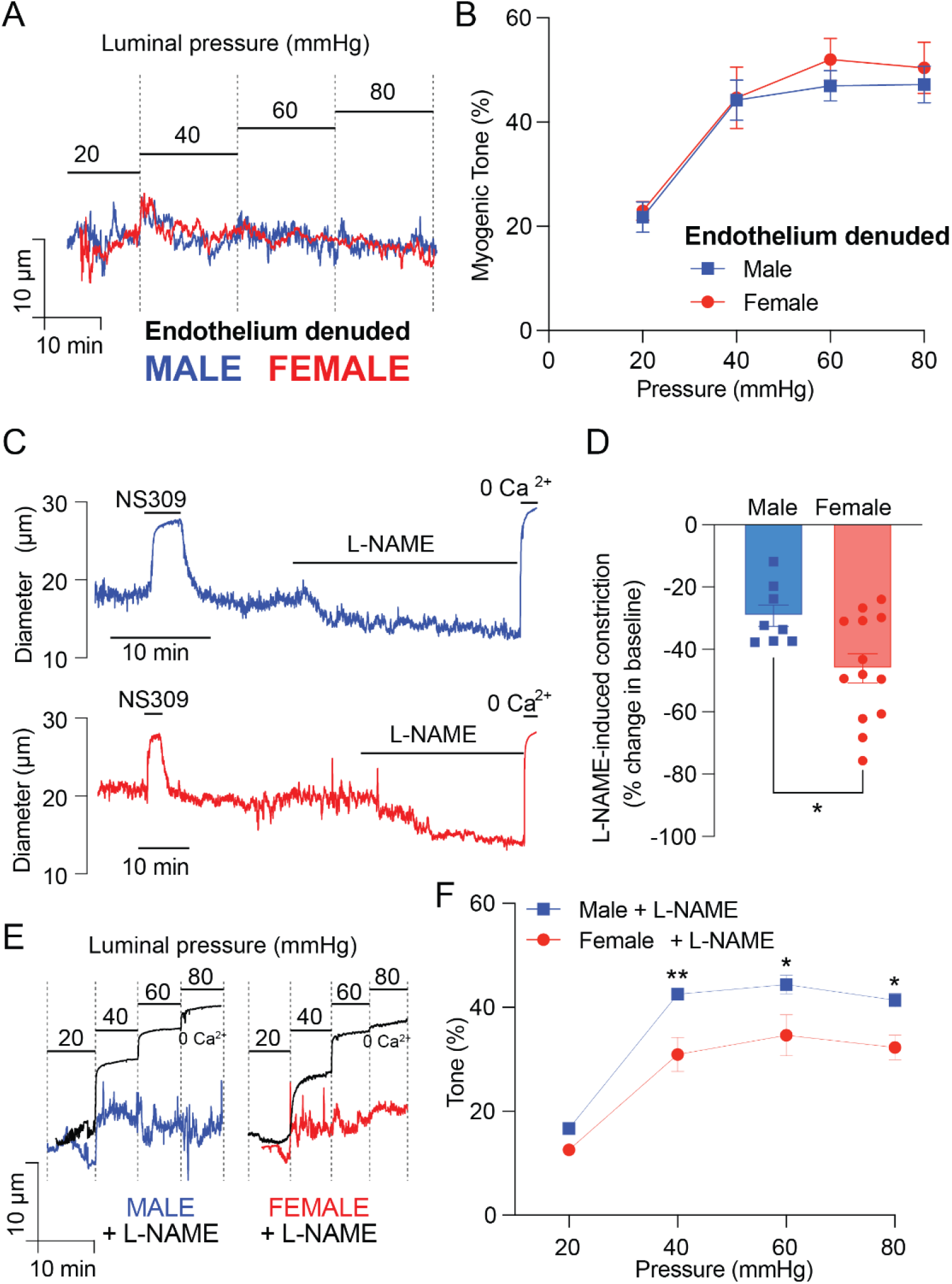
Increased nitric oxide production only partially accounts for the diminished myogenic tone in hiPAs from female mice. (A) Representative endothelium denuded lumen diameter traces from pressure curve myography experiments and (B) plotted myogenic tone in endothelium denuded HiPAs from male and female mice. (C) Representative male and female pressure myography traces showing L-NAME-induced constriction. (D) Comparison of L-NAME induced change in baseline diameter between male and female mice. (E) Representative pressure curve traces showing lumen diameter in the presence of vascular NO production inhibitor L-NAME plotted with passive diameter in 0 Ca2+ aCSF. (F) Comparison of myogenic tone levels over range of pressures between male and female mice in the presence of L-NAME.

The canonical vasodilatory pathway of NO on cerebral SMCs is the activation of soluble guanylyl cyclase (sGC), which increases levels of cyclic guanosine monophosphate (cGMP), activating protein kinase G (PKG), and subsequent opening of large-conductance Ca^2+^-activated K^+^ (BK) channels^40-43^. We then further compared the BK channels activity between male and female arteries by using its blocker paxilline. Consistent with previous observations from intracerebral arterioles^17,44^, 1 μM paxilline had only a modest contractile effect on male HiPAs (Supp Fig. 1B), which was significantly lower (11.52% ± 1.84) compared to the constriction induced in female HiPAs (34.87% ± 5.01). Remarkably, the higher BK channel activity in female arterioles revealed by the paxilline-induced contraction was abolished by pretreatment with L-NAME (Supp Fig. 1B), indicating that this sex difference was driven by the eNOS activity. Decisively, we tested the myogenic responses of male and female HiPAs in presence of L-NAME. While the inhibition of eNOS alone reduced the difference in the constriction induced by 20 mmHg-step increases in luminal pressure, it failed to bring the tone of female HiPAs to the levels measured in males (Fig 3E-F). Taken together, these results indicate that an additional endothelial-dependent mechanism is involved in sex differences.

### Endothelium-Derived Hyperpolarization Is More Pronounced in Female Hippocampal Arterioles

In tandem with the NO release, the endothelium exerts a tonic dilatory influence on intracerebral arterioles via the activity of SK and IK channels^24^. Thus, one potential explanation for the decreased myogenic response observed in female HiPAs could be an increased SK and/or IK channel activity. We then measured the constriction induced by inhibition of SK channels with 100 nM apamin and IK channels with 300 nM charybdotoxin. As charybdotoxin also blocks BK channels, the experiments were conducted in presence of paxilline to narrow down the effect to IK channels only. Interestingly, apamin caused a significantly greater constriction in female HiPAs compared to male ones (22.02 ± 1.74% vs 11.35 ± 1.88%), while charybdotoxin induced similar effects (Fig. 4A-B). Simultaneous addition of apamin and charybdotoxin replicated the difference (Fig. 4B). Finally, we measured myogenic responses of male and female HiPAs in the presence of 100 μM L-NAME and 300 nM apamin and observed virtually no difference between the groups (Fig. 4C -D). Collectively, these data indicate that the lower pressure-induced constriction in female arterioles is mediated by a higher basal NO release and SK channel activity.

**Figure 4:**
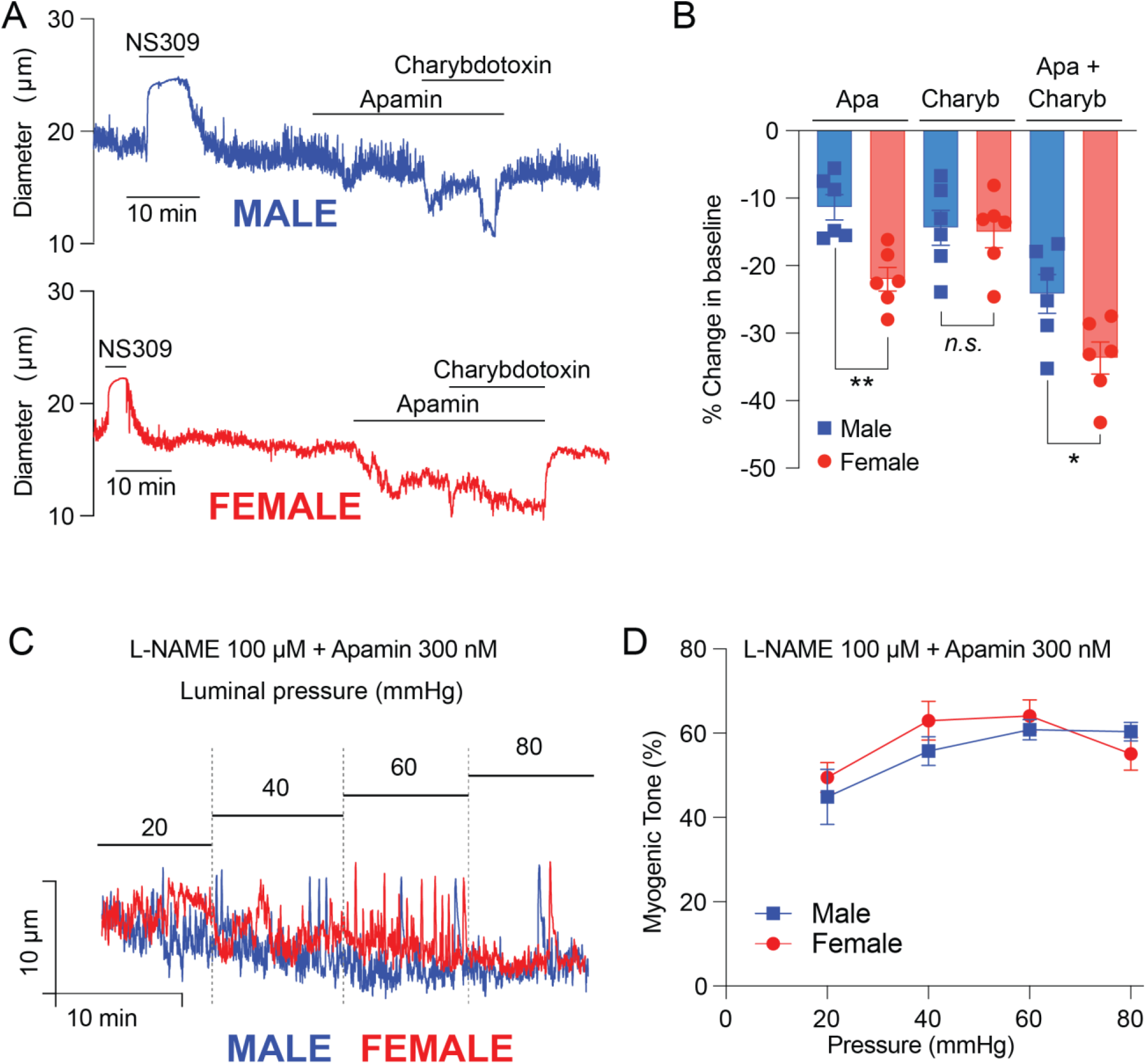
SK channel, but not IK channel, activity is higher in female hippocampal arterioles. (A) Pressure myography traces showing sequential addition of Apamin, SK channel blocker, and Charybdotoxin, IK channel blocker, decrease lumen diameter in both male and female HiPAs. (B) Plotted % change in baseline diameter in male and female mice. (C) Pressure myography curve experiment lumen diameter traces and (D) plotted myogenic tone levels in presence of 100 μM L-NAME and 300nM Apamin. Apamin and Charybdotoxin was used in conjunction with Paxilline to isolate the effects of SK/IK channel reactivity.

### SK Channel Expression Is Increased in Female Hippocampal Microvasculature Compared To Males

Due to the increased SK channel function in female HiPAs, we investigated SK and IK channels expression in hippocampal endothelial cells. Figure 5A displays our novel immunohistochemistry approach to probe protein expression in isolated hippocampal microvasculature using confocal microscopy and Z-stack acquisition. We then developed an unbiased computational analytic workflow to compare IK and SK channel expression in male in female HiPA endothelium (Supp. Fig 2). We found that IK channel protein expression did not differ (2837 ± 599 μm^3^ versus 3062 ± 445 μm3) in male and female endothelial cells, respectfully (Fig. 5B-C). However, and consistent with the enhanced sensitivity towards apamin, SK staining appeared significantly higher in female endothelium (3185 ± 610 μm^3^ of volume versus 1310 ± 373 μm^3^, n = 3 mice). These data indicate that, consistent with the *ex vivo* functional experiments, SK channel expression is increased in hippocampal female endothelial cells compared to male.

**Figure 5:**
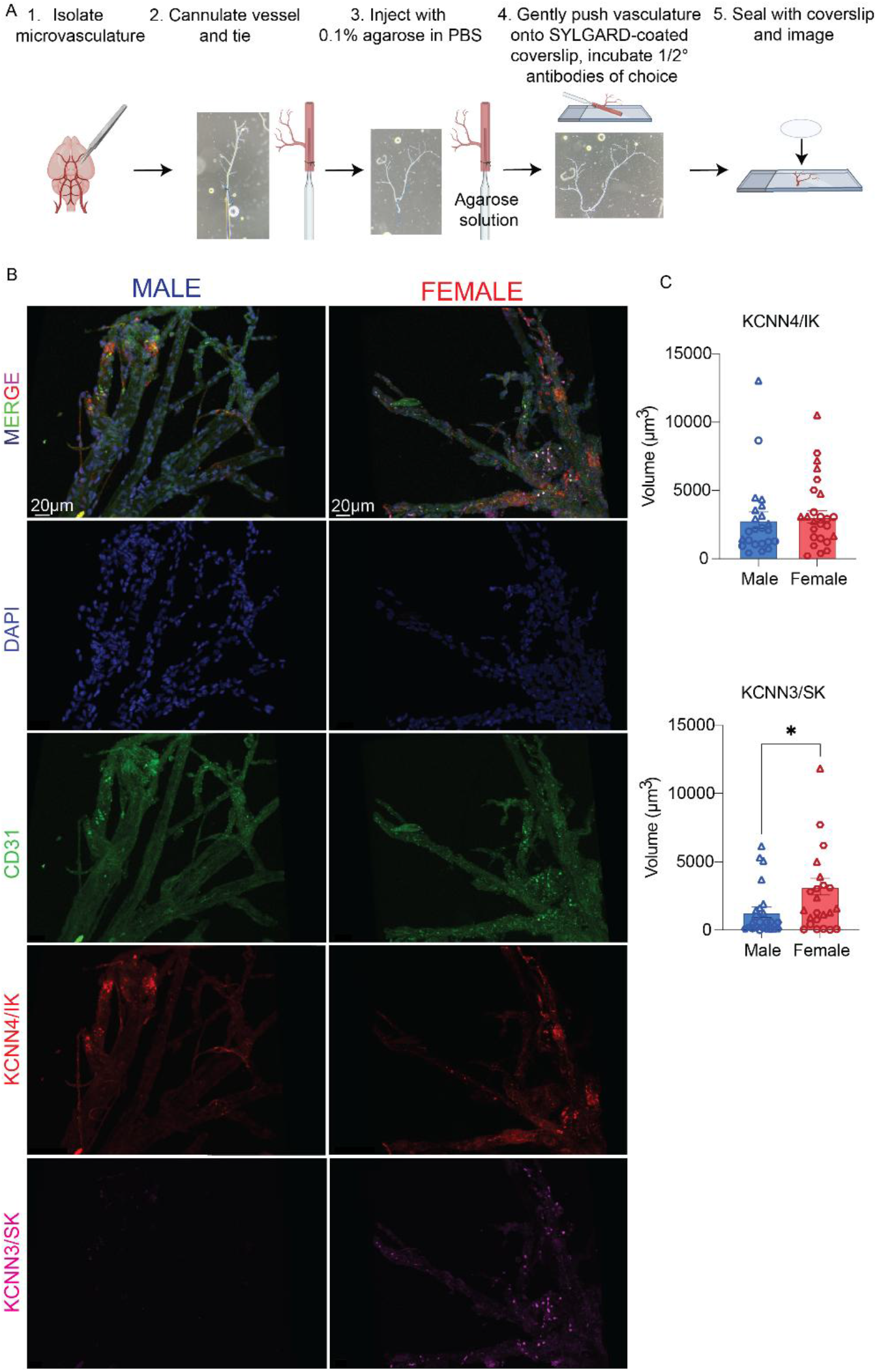
SK channel, but not IK channel, expression is higher in female hippocampal arterioles. (A) Workflow of novel immunohistochemistry staining on isolated microvasculature from the hippocampus. (B) Representative images from male and female hippocampal arterioles. Endothelial cells indicated by CD31 (green), and KCNN4 is IK (red), while KCNN3 is SK (magenta). The merge channel is also indicated. The scale bar is noted and microvascular structures are also indicated. (C) Images were blinded and uploaded into Imaris and volume of fluorescence was taken for both SK and IK expression. 9 ROIs were taken per mouse while symbols correspond to ROIs taken from the same mouse in different vascular regions and vascular segments from isolated hippocampal arterioles, n = 3. * Indicates significance p < 0.05, p = 0.015 via unpaired t-test.

### Circulating Estrogen Underlies Enhanced eNOS and SK Channel Activity in Female HiPAs

The modulatory effect of estrogen on eNOS expression and activity has been well established^8,45^ and more recent studies have suggested a determinant role for estrogen in SK channel gene expression^28,46^. We then explored the impact of ovariectomy (OVX) on L-NAME- and apamin-induced constriction in female HiPAs. Our experimental schematic is shown in Fig. 6A. Three weeks post-surgery, L-NAME induced a 34.57 ± 5.2% constriction of HiPAs from OVX females (n = 5 mice), which was not statistically different from values (31.2 ± 5.36%) obtained in male HiPAs (Fig. 6B). Similarly, OVX brought the apamin-induced constriction to that of males, which was not statistically significant (Fig. 6C, 11.79 ± 2.74 % in males and 17.09 % ± 2.97 % in OVX females). To confirm the central role of estrogen in the decreased myogenic responses from female brain arterioles, we added SC silastic implants to OVX females containing either estrogen or vehicle. We found that OVX females with a vehicle implant had an increased response (constriction) to increases in pressure that was comparable to males (Fig. 6E). Additionally, when OVX mice were given estrogen via implant, the increased response to pressure was ablated (Fig 6 D-E) and returned to female myogenic tone response values seen in Figure 1. Together these results indicate that estrogen enhances eNOS and SK channel activity in arteriolar endothelial cells, resulting in higher retro-control and decreased myogenic tone in female brain parenchymal arterioles compared to males (Fig 7).

**Figure 6:**
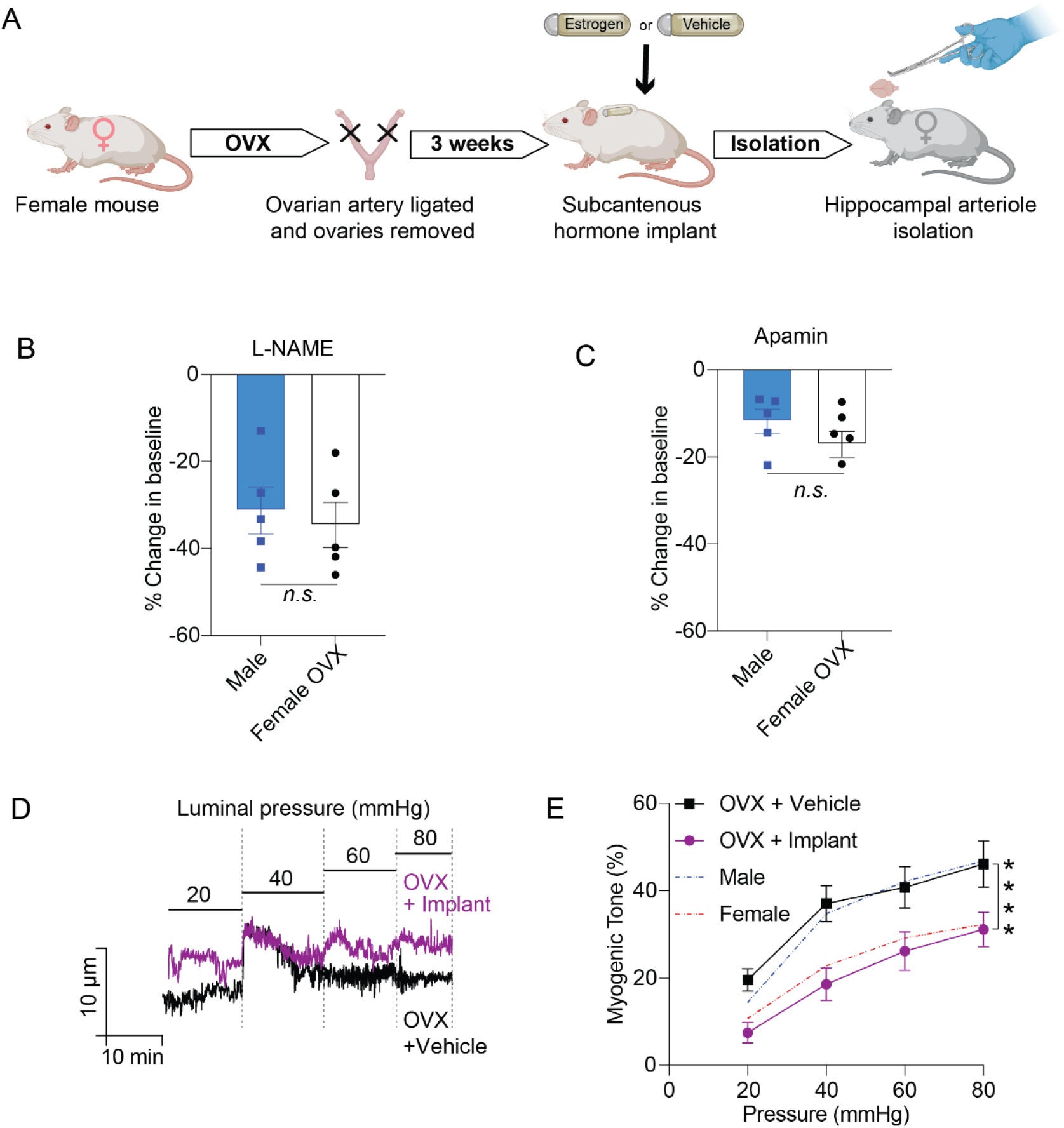
OVX female mice exhibit myogenic tone at levels equivalent to males in HiPAs. Tone levels are restored to physiologically unaltered normal levels in OVX + implant mice. (A) Schematic of ovariectomized (OVX) females with subsequent subcutaneous +/-hormone replacement via silastic implant. Vasculature was removed and mounted using pressure myography interrogation. (B) Change in baseline of male and female OVX HiPAs treated with L-NAME, n = 5 mice for each group, data was not significant via unpaired t-test. (C) Change in baseline percentage for male and female OVX HiPAs treated with Apamin, n = 5 for each group, data was not statistically significant via unpaired t-test. (D) Comparison of female HiPA pressure myography traces over range of physiological pressures in OVX + implant (vehicle or estrogen) mice at 20 mmHg increments over physiologically normal range. (E) Pressure curve plot of myogenic tone percentages over 20 – 80 mmHg in both OVX + vehicle implant and OVX + estrogen implant mice compared to normal myogenic tone levels in physiologically unaltered female and male mice (Figure 1). N = 6 and 9 respectively, **** indicates p < 0.0001 post hoc via Sidak’s multiple comparison test.

**Figure 7:**
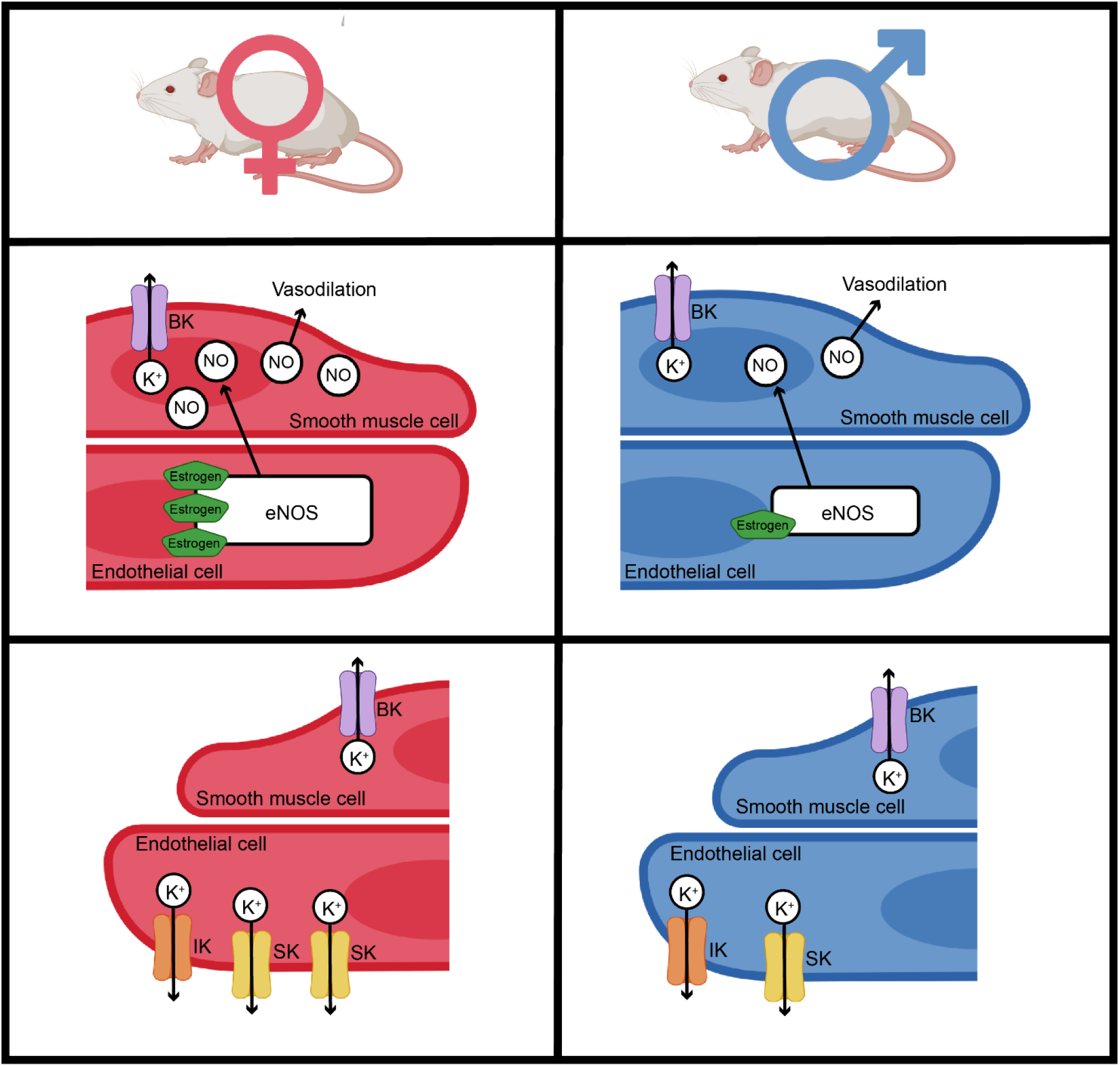
Proposed mechanism: Estrogen regulates myogenic tone in female hippocampal arterioles through release of nitric oxide and increased endothelial cell SK channel activity and expression. Females have decreased myogenic tone due to increased nitric oxide production post estrogen binding to its receptor in endothelial cells. This then causes eNOS activation and resulting NO production, communication with neighboring smooth muscle cells resulting in a robust vasodilation in hippocampal arterioles. Additionally, estrogen causes an upregulation of SK channel expression which we propose then causes increased endothelial-induced hyperpolarization due to K^+^ efflux and further vasodilation. In females this estrogen-dependent response produces regenerative hyperpolarization that goes to neighboring endothelial cells that results in a more pronounced vasodilation and decreased myogenic tone compared to their male counterparts.

## Discussion

The regulation of cerebral blood flow is vital for proper brain function, and it has been suggested that there may be sex-dependent differences in this regulation throughout the brain. Studies considering sex as a biological factor have revealed differences in cerebral arteries in rats where Geary et. al., found that estrogen caused a reduction in female myogenic tone when compared to males^8^. Myogenic tone was originally described by William Bayliss in 1902, that refers to the intrinsic contractile activity of smooth muscle cells in blood vessels in the absence of external hormonal or neuronal influences^11^. Myogenic tone is a fundamental property of the vascular system that plays a crucial role in the regulation of blood pressure and flow. This is carried out by smooth muscle cells in which they exhibit spontaneous rhythmic contractions and relaxation, generating a basal level of tone in the vascular wall^47,48^. Importantly, this phenomenon allows blood vessels to respond dynamically to changes in blood pressure and maintain vital perfusion to systematic organs^11,47-49^. In this study we confirm differences in myogenic tone *in vivo* in the cerebral cortex arterioles while also focusing on a specific brain region, the hippocampus. The hippocampus, a region of the brain known for its crucial role in memory and spatial navigation, receives blood supply through a network of arterioles^50^. Using both male and female C57 age-matched mice, we highlight the differences in male and female brain hippocampal arterioles (HiPAs) where we found that females exhibit lower myogenic tone when compared to their male counterparts.

We initially found lower myogenic tone responses in response to aCSF depleted of Ca^2+^ administered over an acute cranial window of female mice compared to males via TPLSM *in vivo* network diameter measurements. We then wanted to hone in on a specific region of the brain and due to the sex differences reported in hippocampal region including cognition and plasticity^50^, we decided to interrogate hippocampal arterioles. Looking from a vascular standpoint, the vascular density in the hippocampus is a fraction of that of the neocortex^51-53^. Interestingly the formation of the hippocampal arterioles perpendicularly penetrates into the coronal plane and due to their more sparse vascular network requires oxygen to diffuse further^54^, which may result in a decreased response with myogenic tone to have more profound consequences in this brain region. Utilizing *ex vivo* pressure myography from isolated HiPAs we found that there was significantly decreased myogenic tone, supporting our *in vivo* findings. To further characterize our observed sexually dimorphic responses, we then looked to the components of the vasculature.

The vasculature is made up of both an inner endothelial cell layer that forms the walls of the blood vessels and the external mural cells, including both smooth muscle cells an pericytes^55^, therefore we wanted to elucidate which cell type could be contributing to our observed myogenic tone differences in males and females. We found that when we denuded the endothelial layer, differences in myogenic tone were ablated, indicating endothelial-derived and dependent differences. While myogenic tone provides a baseline of vascular constriction, it can be influenced by various neural, hormonal, and metabolic factors such as endothelium-derive nitric oxide (NO)^8,46,56^. Therefore, we investigated if nitric oxide was contributing to these differences. Others have found that the hippocampal vasculature is unique in male rats regarding NO function and that hippocampal arterioles do not respond to L-NAME, however was able to dilate with NO-dependent PAI-1 inhibition, suggesting eNOS was present and expression in their vasculature^57^. We however found that both males and female HiPAs constricted to the nitric oxide inhibitor, L-NAME, whereas females had a significantly larger constriction. Differences in our L-NAME-induced vasoreactivity could be due Johnson et. al.’s isolation of hippocampal arterials supplying the CA1 in the dorsal hippocampus^57^ while we surveyed all vasculature from the entire hippocampus. Importantly in our studies, L-NAME only partially reduced these differences, indicating increased NO production in females was not the only player contributing to decreased myogenic tone. Large-conductance, voltage-dependent BK_Ca_ channels also play a major role in myogenic tone in pial arteries and are highly expressed by arteriolar smooth muscle cells^24,58^. We were curious if this channel also exhibited sexual dimorphism. Our results show increased BK_Ca_-dependent constriction upon Paxilline (BK_Ca_ channel blocker) administration in female HiPAs, however when we inhibited eNOS production via L-NAME these differences were ablated, further solidifying an endothelial cell-induced change in female arteriolar diameter size.

Due to the established role of endothelial small- and intermediate-conductance calcium-sensitive K+ channels (SK_Ca_ and IK_Ca_ channels) in regulating brain parenchymal arterial diameter and corresponding cerebral blood flow^23,24,59^, we examined endothelium-derived hyperpolarization. SK channels are Ca^2+^ sensitive and are not voltage-dependent and are responsible for regulating the flow of K^+^ ions across the cell membrane^24^. They are involved in the generation of feedback slow after-hyperpolarization via efflux of K^+^ which leads to inhibition of setting an action potential and thus causes vascular relaxation^46^. Therefore, to elucidate the role of this channel in our observed myogenic tone changes we utilized a pharmaceutical approach to probe the various channels expressed in arteriole endothelial cells to interrogate potential players. We found that inhibition of SK channels using Apamin caused a greater constriction in female HiPAs, which was abolished by pretreatment with the NO inhibitor. Furthermore, we found that there was increased expression of SK channels in female HiPAs compared to males, which suggests that the differences in expression may account for observed functional changes. SK channels are also significant regulators of afterhyperpolarization in neurons to counterbalance action potential frequency^60-62^; thus, it would be interesting to see if this increased SK function and expression in females extends into the neuronal network in the hippocampus. Taken together our findings suggest that eNOS and SK channels have a greater influence on vasodilation and myogenic tone in female mice at physiological pressures, indicating a sex difference in the regulation of these mechanisms in the hippocampus.

The hippocampus contains several sex hormone receptors including androgen and estrogen receptors (α, β, and G-coupled protein receptors)^50^. We hypothesize that sex-linked hormone regulation of myogenic tone also affects the microcirculation in the hippocampus, therefore we wanted to look at how estrogen plays a role. Estrogen receptors in the hippocampus have been shown to be sexually dimorphic, where others have found a more significant number of neuronal ER-β receptors in female mice compared to males^63^, however this was not explored in the vasculature. Estrogen regulates myogenic tone by rapidly inducing endothelial nitric oxide synthase (eNOS) which promotes nitric oxide (NO) release which results in vasodilation^8,64,65^. Research has shown that estrogen lowers myogenic tone in female mice by enhancing the release of nitric oxide from the endothelium in the middle cerebral artery and corresponding parenchymal arterioles^8,66^, however it remains unclear whether this difference extends to the microcirculation within the hippocampus. Additionally, previous work has shown estrogen causes increased endothelial cell SK expression and function in colonic smooth muscle cells^46^, therefore we wanted to investigate this pipeline in HiPAs. To reduce estrogen in our female cohort, an ovariectomy (OVX) was performed in female mice and HiPAs were isolated three weeks later. We found the Vaso-responses to NO inhibitor and SK channel blocker in our OVX female mice decreased to levels like those seen in males. These results support that estrogen influences our reported unique increased response to L-NAME as well as increased function and probable expression of SK channels. Furthermore, OVX females had increased myogenic tone in physiological pressures which equalized to that of males. When estrogen was administered to OVX females, the responses to increases in pressure measuring myogenic tone returned to the baseline levels observed in intact females. These findings further underscore the prominence of estrogen in regulating hippocampal blood flow and possibly leading to differences in male and females in neurodegenerative disease progression.

There is ongoing research exploring the impact of hormone replacement therapy (HRT) on neurodegenerative diseases such as Alzheimer’s disease (AD) or Parkinson’s Disease. In AD, women show increased disease prevalence as well as more severe cognitive symptoms while in schizophrenia and Parkinson’s Disease men show more severe cognitive impairment^50,67^. Therefore, understanding sexual dimorphism in the microcirculation will contribute to better targeted patient-specific therapies in neurodegenerative diseases. Estrogen is particularly of interest in AD due to its potential neuroprotective effects^68^. Three main estrogens are physiologically synthesized during the reproductive lifetime of women including estrone (E1), 17β-estradiol (E2), and estriol (E3), where postmenopausal the circulating levels fall 1000 fold^69^. Due to the increase in female disease progression postmenopausal and inverse correlation of circulating estrogen, others have studied how estrogen treatment could reduce AD disease progression. Early studies were promising showing that E2 HRT in an AD model prevented neuronal loss in OVX rats^70^. While estrogen has been reported to have protective effects for cerebral endothelial cells, HRT has not had success in recent randomized clinical trials looking delaying the onset of AD^71^. Thus, HRT of estrogen may not be the solution perhaps because there are changes in estrogen receptors postmenopausal. Additionally other factors such as lifestyle factors, genetic risk, and timing may play a role in why the HRT clinical trials have been inconsistent at best^71^. Our findings suggest sex differences in the responsiveness of hippocampal arterioles to various physiological and pathological stimuli which contributes to divergent outcomes when regulating appropriate hippocampal blood flow. This decrease in myogenic tone observed in female mice may result in neuronal-energetic deficits and a resulting decrease in cognitive function and sex-dependent susceptibility to neurological disorders.

Sex differences in hippocampal arterioles represent an important area of research in neuroscience. Variations in size, morphology, and functionality of these blood vessels between males and females may contribute to differences in cognitive function and susceptibility to neurological disorders. Our work provides evidence for sex differences in the reactivity of hippocampal arterioles in mice. These differences are influenced by endothelium-dependent mechanisms, including nitric oxide and endothelium-derived hyperpolarization mediated by SK channels, contributing to decreased myogenic tone in females compared to males. Further investigations into the underlying mechanisms, estrogen-specific receptors, and implications of these sex differences can contribute to our understanding of brain function and potentially lead to sex-specific therapeutic interventions in the future.

## Author contributions

DAJ, AR, MBG, PR, JTF, ACR and FD performed *ex vivo* experiments, data collection, and analysis. DAJ performed all *in vivo* experiments, data collection and analysis. DAJ and AR performed the IHC, data collection and analysis. ACR performed the OVX surgeries. DAJ and FD designed and directed the research as well as wrote the manuscript. All authors edited the manuscript and approved its submission.

## Acknowledgement

This study was supported by a research grant from the Center for Women’s Health Research located at the University of Colorado Anschutz Medical Campus; 2 research grants from the University of Pennsylvania Orphan Disease Center in partnership with the cureCADASIL (2019 and 2022), the National Institute of Neurological Disorders and Stroke RF1/R01NS129022, and the National Heart, Lung, and Blood Institute R01HL136636 to FD and 5T32GM007635, and National Heart, Lung, and Blood Institute F31HL170645 to DAJ.

## Conflict of interest

The authors declare no competing financial interests.

**Supplemental Figure 1:**
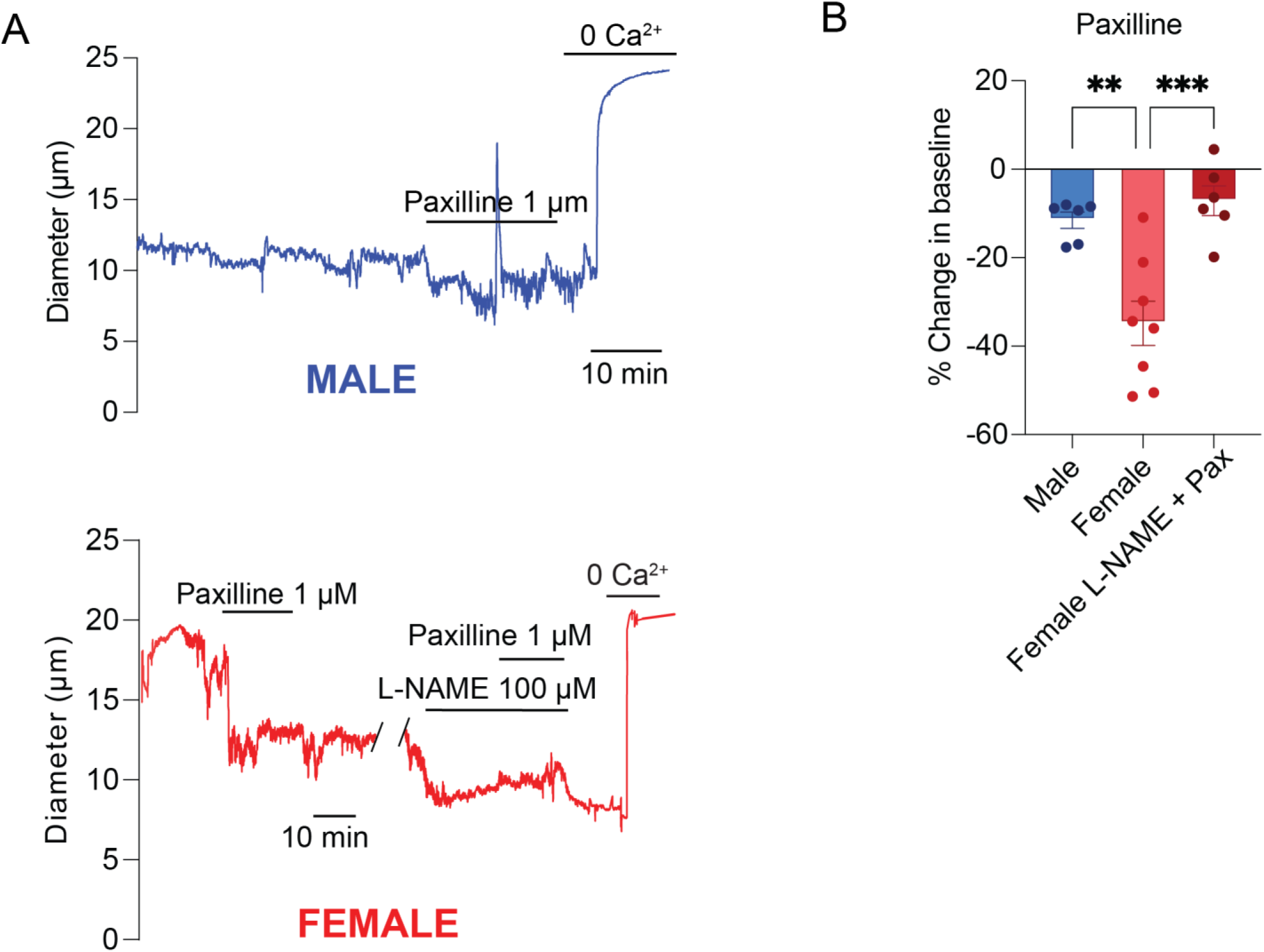
BK channel activity is increased in female hippocampal arterioles while eNOS inhibition abolished this increased activity. (A) Representative pressure myography traces of male treated with Paxilline and females with both Paxilline and Paxilline + L-NAME. Concentration and duration are noted on each trace. (B) Change in baseline for female and male treated HiPAs with 1 μM Paxilline. N = 5, 8, and 6 for males, females, and females treated with L-NAME and Paxilline, respectively. ** indicates p < 0.01, *** indicates p < 0.001. Error bars represent ± SEM.

**Supplemental Figure 2:**
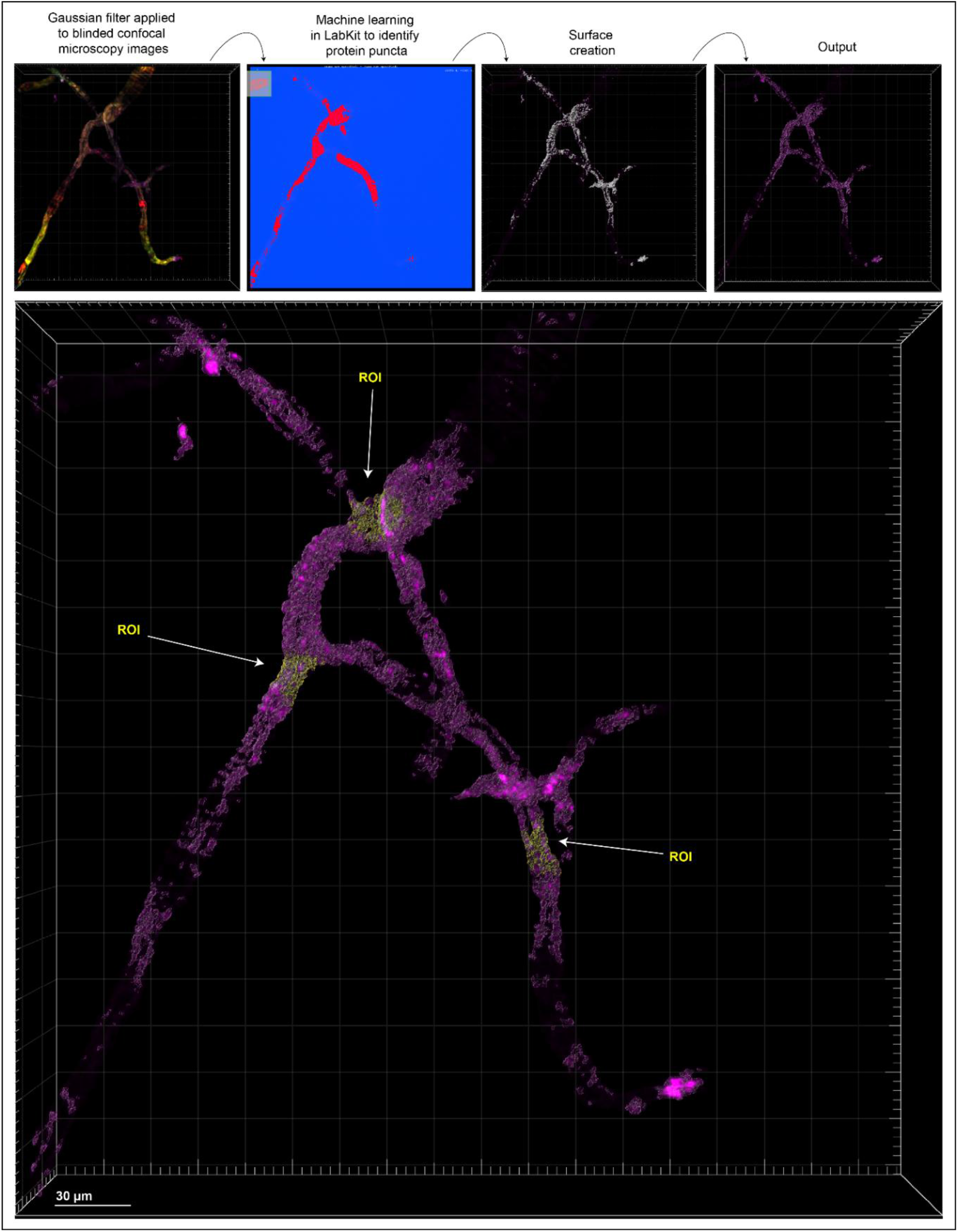
Imaris workflow for immunohistochemistry analysis. Blinded images were uploaded into Imaris and a gaussian filter was applied to reduce background noise. Fluorescent puncta surface creation was implemented using LabKit where the same classifier was used for all images for each respective antibody. Then ROIs were randomly selected from each individual mouse and total fluorescent volume from each Z-stack was collected.

## References

1 Regitz-Zagrosek, V. & Kararigas, G. Mechanistic Pathways of Sex Differences in Cardiovascular Disease. Physiol Rev 97, 1–37, doi:10.1152/physrev.00021.2015 (2017).

2 Mendelsohn, M. E. & Karas, R. H. The protective effects of estrogen on the cardiovascular system. N Engl J Med 340, 1801–1811, doi:10.1056/NEJM199906103402306 (1999).

3 Mendelsohn, M. E. & Karas, R. H. Molecular and cellular basis of cardiovascular gender differences. Science 308, 1583–1587, doi:10.1126/science.1112062 (2005).

4 Forouzanfar, M. H. et al. Global Burden of Hypertension and Systolic Blood Pressure of at Least 110 to 115 mm Hg, 1990-2015. JAMA 317, 165–182, doi:10.1001/jama.2016.19043 (2017).

5 Wellman, G. C., Bonev, A. D., Nelson, M. T. & Brayden, J. E. Gender differences in coronary artery diameter involve estrogen, nitric oxide, and Ca(2+)-dependent K+ channels. Circ Res 79, 1024–1030, doi:10.1161/01.res.79.5.1024 (1996).

6 Lamping, K. G. & Faraci, F. M. Role of sex differences and effects of endothelial NO synthase deficiency in responses of carotid arteries to serotonin. Arterioscler Thromb Vasc Biol 21, 523–528, doi:10.1161/01.atv.21.4.523 (2001).

7 Skarsgard, P., van Breemen, C. & Laher, I. Estrogen regulates myogenic tone in pressurized cerebral arteries by enhanced basal release of nitric oxide. Am J Physiol 273, H2248–2256, doi:10.1152/ajpheart.1997.273.5.H2248 (1997).

8 Geary, G. G., Krause, D. N. & Duckles, S. P. Estrogen reduces myogenic tone through a nitric oxide-dependent mechanism in rat cerebral arteries. Am J Physiol 275, H292–300, doi:10.1152/ajpheart.1998.275.1.H292 (1998).

9 Lundberg, J. O., Gladwin, M. T. & Weitzberg, E. Strategies to increase nitric oxide signalling in cardiovascular disease. Nat Rev Drug Discov 14, 623–641, doi:10.1038/nrd4623 (2015).

10 Vanhoutte, P. M., Shimokawa, H., Feletou, M. & Tang, E. H. Endothelial dysfunction and vascular disease - a 30th anniversary update. Acta Physiol (Oxf) 219, 22–96, doi:10.1111/apha.12646 (2017).

11 Bayliss, W. M. On the local reactions of the arterial wall to changes of internal pressure. J Physiol 28, 220–231, doi:10.1113/jphysiol.1902.sp000911 (1902).

12 Faraci, F. M. & Heistad, D. D. Regulation of large cerebral arteries and cerebral microvascular pressure. Circ Res 66, 8–17, doi:10.1161/01.res.66.1.8 (1990).

13 Claassen, J., Thijssen, D. H. J., Panerai, R. B. & Faraci, F. M. Regulation of cerebral blood flow in humans: physiology and clinical implications of autoregulation. Physiol Rev 101, 1487–1559, doi:10.1152/physrev.00022.2020 (2021).

14 Knot, H. J. & Nelson, M. T. Regulation of membrane potential and diameter by voltage-dependent K+ channels in rabbit myogenic cerebral arteries. Am J Physiol 269, H348–355, doi:10.1152/ajpheart.1995.269.1.H348 (1995).

15 Knot, H. J., Standen, N. B. & Nelson, M. T. Ryanodine receptors regulate arterial diameter and wall [Ca2+] in cerebral arteries of rat via Ca2+-dependent K+ channels. J Physiol 508 (Pt 1), 211–221, doi:10.1111/j.1469-7793.1998.211br.x (1998).

16 Brian, J. E., Jr., Heistad, D. D. & Faraci, F. M. Effect of carbon monoxide on rabbit cerebral arteries. Stroke 25, 639–643; discussion 643-634, doi:10.1161/01.str.25.3.639 (1994).

17 Dabertrand, F. et al. Potassium channelopathy-like defect underlies early-stage cerebrovascular dysfunction in a genetic model of small vessel disease. Proc Natl Acad Sci U S A 112, E796–805, doi:10.1073/pnas.1420765112 (2015).

18 Stanhewicz, A. E., Wenner, M. M. & Stachenfeld, N. S. Sex differences in endothelial function important to vascular health and overall cardiovascular disease risk across the lifespan. Am J Physiol Heart Circ Physiol 315, H1569–H1588, doi:10.1152/ajpheart.00396.2018 (2018).

19 Iorga, A. et al. The protective role of estrogen and estrogen receptors in cardiovascular disease and the controversial use of estrogen therapy. Biol Sex Differ 8, 33, doi:10.1186/s13293-017-0152-8 (2017).

20 Sandow, S. L., Senadheera, S., Bertrand, P. P., Murphy, T. V. & Tare, M. Myoendothelial contacts, gap junctions, and microdomains: anatomical links to function? Microcirculation 19, 403–415, doi:10.1111/j.1549-8719.2011.00146.x (2012).

21 Andresen, J., Shafi, N. I. & Bryan, R. M., Jr. Endothelial influences on cerebrovascular tone. J Appl Physiol (1985) 100, 318–327, doi:10.1152/japplphysiol.00937.2005 (2006).

22 Edwards, G., Feletou, M. & Weston, A. H. Endothelium-derived hyperpolarising factors and associated pathways: a synopsis. Pflugers Arch 459, 863–879, doi:10.1007/s00424-010-0817-1 (2010).

23 Cipolla, M. J., Smith, J., Kohlmeyer, M. M. & Godfrey, J. A. SKCa and IKCa Channels, myogenic tone, and vasodilator responses in middle cerebral arteries and parenchymal arterioles: effect of ischemia and reperfusion. Stroke 40, 1451–1457, doi:10.1161/STROKEAHA.108.535435 (2009).

24 Hannah, R. M., Dunn, K. M., Bonev, A. D. & Nelson, M. T. Endothelial SK(Ca) and IK(Ca) channels regulate brain parenchymal arteriolar diameter and cortical cerebral blood flow. J Cereb Blood Flow Metab 31, 1175–1186, doi:10.1038/jcbfm.2010.214 (2011).

25 Ledoux, J. et al. Functional architecture of inositol 1,4,5-trisphosphate signaling in restricted spaces of myoendothelial projections. Proc Natl Acad Sci U S A 105, 9627–9632, doi:10.1073/pnas.0801963105 (2008).

26 Si, H. et al. Impaired endothelium-derived hyperpolarizing factor-mediated dilations and increased blood pressure in mice deficient of the intermediate-conductance Ca2+-activated K+ channel. Circ Res 99, 537–544, doi:10.1161/01.RES.0000238377.08219.0c (2006).

27 Taylor, M. S. et al. Altered expression of small-conductance Ca2+-activated K+ (SK3) channels modulates arterial tone and blood pressure. Circ Res 93, 124–131, doi:10.1161/01.RES.0000081980.63146.69 (2003).

28 Jacobson, D., Pribnow, D., Herson, P. S., Maylie, J. & Adelman, J. P. Determinants contributing to estrogen-regulated expression of SK3. Biochem Biophys Res Commun 303, 660–668, doi:10.1016/s0006-291x(03)00408-x (2003).

29 Yap, F. C., Taylor, M. S. & Lin, M. T. Ovariectomy-induced reductions in endothelial SK3 channel activity and endothelium-dependent vasorelaxation in murine mesenteric arteries. PLoS One 9, e104686, doi:10.1371/journal.pone.0104686 (2014).

30 Nishimura, N., Schaffer, C. B., Friedman, B., Lyden, P. D. & Kleinfeld, D. Penetrating arterioles are a bottleneck in the perfusion of neocortex. Proc Natl Acad Sci U S A 104, 365–370, doi:10.1073/pnas.0609551104 (2007).

31 Fernandez-Klett, F., Offenhauser, N., Dirnagl, U., Priller, J. & Lindauer, U. Pericytes in capillaries are contractile in vivo, but arterioles mediate functional hyperemia in the mouse brain. Proc Natl Acad Sci U S A 107, 22290–22295, doi:10.1073/pnas.1011321108 (2010).

32 Longden, T. A. et al. Capillary K(+)-sensing initiates retrograde hyperpolarization to increase local cerebral blood flow. Nat Neurosci 20, 717–726, doi:10.1038/nn.4533 (2017).

33 Iadecola, C. The pathobiology of vascular dementia. Neuron 80, 844–866, doi:10.1016/j.neuron.2013.10.008 (2013).

34 Rosehart, A. C., Johnson, A. C. & Dabertrand, F. Ex Vivo Pressurized Hippocampal Capillary-Parenchymal Arteriole Preparation for Functional Study. J Vis Exp, doi:10.3791/60676 (2019).

35 Pires, P. W., Dabertrand, F. & Earley, S. Isolation and Cannulation of Cerebral Parenchymal Arterioles. J Vis Exp, doi:10.3791/53835 (2016).

36 Jeffrey, D., Fontaine, J. T. & Dabertrand, F. Ex vivo capillary-parenchymal arteriole approach to study brain pericyte physiology. Neurophotonics 9, 031919 (2022).

37 Ralevic, V., Kristek, F., Hudlicka, O. & Burnstock, G. A new protocol for removal of the endothelium from the perfused rat hind-limb preparation. Circ Res 64, 1190–1196, doi:10.1161/01.res.64.6.1190 (1989).

38 Bird, C. M. & Burgess, N. The hippocampus and memory: insights from spatial processing. Nat Rev Neurosci 9, 182–194, doi:10.1038/nrn2335 (2008).

39 Baumbach, G. L., Sigmund, C. D. & Faraci, F. M. Cerebral arteriolar structure in mice overexpressing human renin and angiotensinogen. Hypertension 41, 50–55, doi:10.1161/01.hyp.0000042427.05390.5c (2003).

40 Robertson, B. E., Schubert, R., Hescheler, J. & Nelson, M. T. cGMP-dependent protein kinase activates Ca-activated K channels in cerebral artery smooth muscle cells. Am J Physiol 265, C299–303, doi:10.1152/ajpcell.1993.265.1.C299 (1993).

41 Sobey, C. G. & Faraci, F. M. Effect of nitric oxide and potassium channel agonists and inhibitors on basilar artery diameter. Am J Physiol 272, H256–262, doi:10.1152/ajpheart.1997.272.1.H256 (1997).

42 Sobey, C. G. & Faraci, F. M. Effects of a novel inhibitor of guanylyl cyclase on dilator responses of mouse cerebral arterioles. Stroke 28, 837–842; discussion 842-833, doi:10.1161/01.str.28.4.837 (1997).

43 Faraci, F. M., Mayhan, W. G., Schmid, P. G. & Heistad, D. D. Effects of arginine vasopressin on cerebral microvascular pressure. Am J Physiol 255, H70–76, doi:10.1152/ajpheart.1988.255.1.H70 (1988).

44 Dabertrand, F., Nelson, M. T. & Brayden, J. E. Acidosis dilates brain parenchymal arterioles by conversion of calcium waves to sparks to activate BK channels. Circ Res 110, 285–294, doi:10.1161/CIRCRESAHA.111.258145 (2012).

45 Zhao, Y., Vanhoutte, P. M. & Leung, S. W. Vascular nitric oxide: Beyond eNOS. J Pharmacol Sci 129, 83–94, doi:10.1016/j.jphs.2015.09.002 (2015).

46 Tang, Y. R. et al. Estrogen regulates the expression of small-conductance Ca-activated K+ channels in colonic smooth muscle cells. Digestion 91, 187–196, doi:10.1159/000371544 (2015).

47 Davis, M. J. & Hill, M. A. Signaling mechanisms underlying the vascular myogenic response. Physiol Rev 79, 387–423, doi:10.1152/physrev.1999.79.2.387 (1999).

48 Jackson, W. F. Myogenic Tone in Peripheral Resistance Arteries and Arterioles: The Pressure Is On! Front Physiol 12, 699517, doi:10.3389/fphys.2021.699517 (2021).

49 Capone, C. et al. Mechanistic insights into a TIMP3-sensitive pathway constitutively engaged in the regulation of cerebral hemodynamics. Elife 5, doi:10.7554/eLife.17536 (2016).

50 Yagi, S. & Galea, L. A. M. Sex differences in hippocampal cognition and neurogenesis. Neuropsychopharmacology 44, 200–213, doi:10.1038/s41386-018-0208-4 (2019).

51 Kirst, C. et al. Mapping the Fine-Scale Organization and Plasticity of the Brain Vasculature. Cell 180, 780–795 e725, doi:10.1016/j.cell.2020.01.028 (2020).

52 Nair, V., Palm, D. & Roth, L. J. Relative vascularity of certain anatomical areas of the brain and other organs of the rat. Nature 188, 497–498, doi:10.1038/188497a0 (1960).

53 Shaw, K. et al. Neurovascular coupling and oxygenation are decreased in hippocampus compared to neocortex because of microvascular differences. Nat Commun 12, 3190, doi:10.1038/s41467-021-23508-y (2021).

54 Ji, X. et al. Brain microvasculature has a common topology with local differences in geometry that match metabolic load. Neuron 109, 1168–1187 e1113, doi:10.1016/j.neuron.2021.02.006 (2021).

55 Daneman, R. & Prat, A. The blood-brain barrier. Cold Spring Harb Perspect Biol 7, a020412, doi:10.1101/cshperspect.a020412 (2015).

56 de Wit, C., Jahrbeck, B., Schafer, C., Bolz, S. S. & Pohl, U. Nitric oxide opposes myogenic pressure responses predominantly in large arterioles in vivo. Hypertension 31, 787–794, doi:10.1161/01.hyp.31.3.787 (1998).

57 Johnson, A. C., Miller, J. E. & Cipolla, M. J. Memory impairment in spontaneously hypertensive rats is associated with hippocampal hypoperfusion and hippocampal vascular dysfunction. J Cereb Blood Flow Metab 40, 845–859, doi:10.1177/0271678X19848510 (2020).

58 Brayden, J. E. & Nelson, M. T. Regulation of arterial tone by activation of calcium-dependent potassium channels. Science 256, 532–535, doi:10.1126/science.1373909 (1992).

59 Pires, P. W., Sullivan, M. N., Pritchard, H. A., Robinson, J. J. & Earley, S. Unitary TRPV3 channel Ca2+ influx events elicit endothelium-dependent dilation of cerebral parenchymal arterioles. Am J Physiol Heart Circ Physiol 309, H2031–2041, doi:10.1152/ajpheart.00140.2015 (2015).

60 Deignan, J. et al. SK2 and SK3 expression differentially affect firing frequency and precision in dopamine neurons. Neuroscience 217, 67–76, doi:10.1016/j.neuroscience.2012.04.053 (2012).

61 Matschke, L. A. et al. Calcium-activated SK potassium channels are key modulators of the pacemaker frequency in locus coeruleus neurons. Mol Cell Neurosci 88, 330–341, doi:10.1016/j.mcn.2018.03.002 (2018).

62 Sun, J., Liu, Y., Baudry, M. & Bi, X. SK2 channel regulation of neuronal excitability, synaptic transmission, and brain rhythmic activity in health and diseases. Biochim Biophys Acta Mol Cell Res 1867, 118834, doi:10.1016/j.bbamcr.2020.118834 (2020).

63 Zhang, J. Q., Cai, W. Q., Zhou, D. S. & Su, B. Y. Distribution and differences of estrogen receptor beta immunoreactivity in the brain of adult male and female rats. Brain Res 935, 73–80, doi:10.1016/s0006-8993(02)02460-5 (2002).

64 Haynes, M. P., Li, L., Russell, K. S. & Bender, J. R. Rapid vascular cell responses to estrogen and membrane receptors. Vascul Pharmacol 38, 99–108, doi:10.1016/s0306-3623(02)00133-7 (2002).

65 Hisamoto, K. & Bender, J. R. Vascular cell signaling by membrane estrogen receptors. Steroids 70, 382–387, doi:10.1016/j.steroids.2005.02.011 (2005).

66 Krause, D. N., Duckles, S. P. & Pelligrino, D. A. Influence of sex steroid hormones on cerebrovascular function. J Appl Physiol (1985) 101, 1252–1261, doi:10.1152/japplphysiol.01095.2005 (2006).

67 Bianco, A., Antonacci, Y. & Liguori, M. Sex and Gender Differences in Neurodegenerative Diseases: Challenges for Therapeutic Opportunities. Int J Mol Sci 24, doi:10.3390/ijms24076354 (2023).

68 Li, X. T. The modulation of potassium channels by estrogens facilitates neuroprotection. Front Cell Dev Biol 10, 998009, doi:10.3389/fcell.2022.998009 (2022).

69 Cui, J., Shen, Y. & Li, R. Estrogen synthesis and signaling pathways during aging: from periphery to brain. Trends Mol Med 19, 197–209, doi:10.1016/j.molmed.2012.12.007 (2013).

70 Simpkins, J. W. et al. Role of estrogen replacement therapy in memory enhancement and the prevention of neuronal loss associated with Alzheimer’s disease. Am J Med 103, 19S–25S, doi:10.1016/s0002-9343(97)00260-x (1997).

71 Boyle, C. P. et al. Estrogen, brain structure, and cognition in postmenopausal women. Hum Brain Mapp 42, 24–35, doi:10.1002/hbm.25200 (2021).

